# *De novo* transcriptome sequence of *Senna tora* provides insights into anthraquinone biosynthesis

**DOI:** 10.1101/837385

**Authors:** Sang-Ho Kang, Woo-Haeng Lee, Chang-Muk Lee, Joon-Soo Sim, So Youn Won, So-Ra Han, Soo-Jin Kwon, Jung Sun Kim, Chang-Kug Kim, Tae-Jin Oh

## Abstract

*Senna tora* is an annual herb with rich source of anthraquinones that have tremendous pharmacological properties. However, there is little mention of genetic information for this species, especially regarding the biosynthetic pathways of anthraquinones. To understand the key genes and regulatory mechanism of anthraquinone biosynthesis pathways, we performed spatial and temporal transcriptome sequencing of *S. tora* using short RNA sequencing (RNA-Seq) and long-read isoform sequencing (Iso-Seq) technologies, and generated two unigene sets composed of 118,635 and 39,364, respectively. A comprehensive functional annotation and classification with multiple public databases identified array of genes involved in major secondary metabolite biosynthesis pathways and important transcription factor (TF) families (MYB, MYB-related, AP2/ERF, C2C2-YABBY, and bHLH). Differential expression analysis indicated that the expression level of genes involved in anthraquinone biosynthetic pathway regulates differently depending on the degree of tissues and seeds development. Furthermore, we identified that the amount of anthraquinone compounds were greater in late seeds than early ones. In conclusion, these results provide a rich resource for understanding the anthraquinone metabolism in *S. tora*.

## Introduction

*Senna tora* (Subfamily, Caesalpiniaceae; and Family, Leguminosae) also known as *Cassia tora*, is an annual xerophytic shrub which grows in the arid zones after the rainy season [1]. This plant is mostly found in India, China, Sri Lanka, Nepal, the Korean peninsula, and other Asian countries. Its name varies in different locales such as Foetid Senna tora, Sickle senna, Wild senna, Coffee pod, Tovara, Chakvad, and Ringworm plant. *S. tora* leaves, seeds, and roots have long been used as food ingredients. It is also valued as a medicinal plant in Ayurveda, commonly used as a depurative, antiperiodic, anthelmintic, liver tonic, hepatic disorders, dyspepsia leprosy, constipation, intermittent fever, cough, bronchitis, ringworm infection, ophthalmic, skin diseases, and others [2, 3]. It has also been used as laxative and a tonic, and is popularly served as a roasted tea throughout Korea and China [4]. The seeds of *S. tora* contain a variety of bioactive anthraquinone substances, including chrysophanol, obtusin, obtusifolin, aurantio-obtusin, chyro-obtusin, obstsifolin, emodin, rubrofusarin, gentibioside, and rhein. Chryophanol is primarily responsible for the plant’s pharmacological properties [5, 6]. *S. tora* mainly contains anthraquinone glycosides and flavonoids [7]. Recently, *S. tora* seed extract (STE) and its active compound aurantio-obtusin have been found to suppress degranulation, histamine production, and reactive oxygen species generation, and also to inhibit the production and mRNA expression of cyclooxygenase 2. STE and aurantio-obtusin also suppressed IgE-mediated FcεRI such as phosphorylation of Syk, protein kinase Cµ, phospholipase Cγ, and extracellular signal-regulated kinases. This suggests that STE and aurantio-obtusin can be beneficial to the treatment of allergy-related diseases [8].

Anthraquinones, secondary metabolites occurring in bacteria, fungi, lichens, and higher plants, seem to originate from a variety of different precursors and pathways. There are two pathways leading to anthraquinone biosynthesis in higher plants: the polyketide pathway and the chorismate/*O*-succinylbenzoic acid pathway. The latter occurs in the plant family Rubiaceae and synthesizes aromatic compounds known for a broad spectrum of bioactivity, such as anticancer, cathartic, anti-inflammatory, anti-microbial, diuretic, vasorelaxing, and phytoestrogen activities, and has recently shown therapeutic potential in autoimmune diabetes [9]. Emodin, physicion, aloe-emodin, and rhein isolated from *S. tora* seed shows antifungal properties against phytopathogenic fungi [10]. Likewise, rhein shows high antibacterial activity towards *Porphyromonas gingivalis* and synergistic antibacterial activity with metronidazole or natural compounds, and the recent studies suggest the immunomodulatory activity of rhein [11–13]. The extract of *S. tora* is found to have hypolipidemic activity, hepatoprotective, and antioxidant effects [2, 14, 15]. Anthraquinones from *S. tora* exhibit significant inhibitory properties against angiotensin-converting enzyme (ACE). Among the various anthraquinones, only anthraquinone glycoside demonstrates marked inhibitory activity against ACE [16].

RNA sequencing (RNA-Seq), a technology that can be used to profile the complete gene space of various organisms due to their high throughput, accuracy, and reproducibility, has accelerated the discovery of new genes or analysis of tissue-specific and functional expression patterns in large, complex genomes like those of plants [17–19]. But in the absence of reference genome information considerable small transcripts hinder the accuracy of the construction of RNA sequencing libraries and the efficiency of functional gene prediction or annotation. Short-length RNA sequencing data limit the creation of a longer, accurate contig assembly, resulting in chimeric contigs and/or low gene annotation [20]. Moreover, small laboratories require high sequencing costs due to the need for long reads and high-depth short read sequences to be accurate in *de novo* assembly. Plants with large genomes pose even more difficult as in, for example, the common soybean crop, which has a genome size of ∼1.1Gb [21]. To improve the comprehensive accuracy of gene prediction, there is a need to introduce a new approach, the “Isoform sequencing (Iso-Seq).” Thanks to its long-read technology, Iso-Seq facilitates identifying new isoforms with a high level of accuracy [22]. Advances in technology enable long reads in the range of 1.5-10 kb, which are able to provide full-length mRNA isoforms, detect new isoforms, and skip the transcript reconstruction process by identifying isoforms directly [23]. In this study, we present the transcriptome analysis of the plant *S. tora* from 4 different sources using RNA-Seq and Iso-Seq, providing insights of key genes involved in anthraquinone biosynthesis in the pharmacologically important herb *S. tora*.

## Materials and methods

### Plant material and RNA preparation

Specimens of *S. tora* (cv. Myeongyun) grown in an experimental plot of National Institute of Horticultural and Herbal Science (Eumseong) field were used for transcriptome analysis. Leaf, root, and early- and late-stage seed tissues were harvested from healthy plants, and stored at −80°C until used for RNA extraction. Total RNA was extracted from leafs, roots, and two stages of seeds of *S. tora* using the RNeasy Plant Mini kit (Qiagen, InS., Valencia, CA, USA). RNA purity was determined using NanoDrop8000 Spectrophotometer and Agilent Technologies 2100 Bioanalyzer, and total RNA integrity was identified as having a minimum integrity value of 7.

### Illumina short-read sequencing

The poly (A)^+^ mRNA was purified and fragmented from 1 µg of total RNA using poly-T oligo-attached magnetic beads by two rounds of purification. Using reverse transcriptase, random hexamer primers, and dUTP, the randomly-cleaved RNA fragments were transcribed reversely into first-strand cDNA. A single A-base was added to these cDNA fragments followed by adapter ligation. The products were purified and concentrated by PCR in order to generate a final-strand specific cDNA library. The quality of the amplified libraries was verified using capillary electrophoresis (Bioanalyzer, Agilent). Quantitative PCR (qPCR) was carried out using SYBR Green PCR Master Mix (Applied Biosystems). Then we pooled together equimolar amounts of libraries that were index-tagged. The cBot-automated cluster creation system (Illumina) performed cluster generation in the flow cell. The sequencing was performed with 2 x 100 bp read length of the flow cell loaded on a HiSeq 2500 sequencing system (Illumina).

### Long-read sequencing

Libraries for Pacific Biosciences Single Molecule Real Time (SMRT) sequencing were prepared from the aforementioned cDNAs. Cycle optimization was performed to determine the optimal number of cycles for large-scale PCR. We prepared 3 fraction cDNAs (1-2 kb, 2-3 kb, and 3-6 kb) using the BluePippin Size selection system. The SMRTbell library was constructed by using SMRTbell^TM^ Template Prep Kit (PN 100-259-100). The DNA/Polymerase Binding Kit P6 (PacBio) was used for DNA synthesis after the sequencing primer annealed to the SMRTbell template. Following the polymerase binding reaction, the MagBead Kit was used to bind the library complex with MagBeads before sequencing. MagBead-bound cDNA complexes result in increased number of reads per SMRT cell. This polymerase-SMRTbell-adaptor complex was then loaded into zero-mode waveguides (ZMWs). The SMRTbell library was sequenced using 8 SMRT cells (Pacific Biosciences) with C4 chemistry (DNA sequencing Reagent 4.0). 1 × 240 minute movies were captured for each SMRT cell using the PacBio RS II sequencing platform.

### De novo transcriptome assembly and sequence clustering

Raw data of the *S. tora* transcriptome generated from Illumina HiSeq were preprocessed to remove nonsense sequences including adaptors, primers, and low quality sequences (Phred quality score of less than 20) using NGS QC Toolkit [24]. The raw data were further processed to remove ribosomal RNA using riboPicker v0.4.3 [25]. The preprocessed reads were then assembled using Trinity [26]. Assembly statistics were calculated using in-house Perl scripts. Assembled transcripts were clustered (CD-HIT-EST v4.6.1) [27] in order to reduce sequence redundancy. Sequence identity threshold and alignment coverage (for the shorter sequence) were both set as 90% to generate clusters. Such clustered transcripts are defined as reference transcripts in this work.

### Illumina expression quantification and differential expression analysis

The cleaned reads from each tissue were aligned with the abundant transcriptome assembly using Bowtie2 [28]. The aligned reads were quantified as fragments per million reads (FPKMs) against non-redundant combined transcript sequences (at 90% sequence similarity by CD-HIT-EST). The reads counting of alignments was performed using RSEM (RNA-Seq by Expectation Maximization)-1.2.25 [29]. The differential expression analysis was performed using the DESeq2 packages [30]. Differentially expressed genes (DEGs) were identified using the combined criteria of a more than twofold change and significance with P-value threshold of 0.001 based on the three biological replicates.

### Functional annotation and classification

All the assembled unigenes were annotated by BLAST program [31] against the National Center for Biotechnology Information (NCBI) nonredundant (Nr) protein database, the Swiss-Prot protein database, and the Kyoto Encyclopedia of Genes and Genomes (KEGG) pathways database with an E-value cutoff of 10^-5^. The best aligning results were selected to annotate the unigenes. Whenever the aligning results from different databases conflicted, the results from Swiss-Prot database were preferentially selected, followed by Nr database and KEGG database. Functional categorization by Geno Ontology (GO) terms [32] was carried out by Blast2GO program [33] with E-value threshold of 10^-5^. AgriGO [34] was used to determine over-representation of GO categories (e.g., biological processes).

### Identification of transcription factor families

To investigate the putative transcription factor families in *S. tora*, unigenes were mapped against all the transcription factor protein sequences made available by the Plant Transcription Factor Database (PlantTFDB 4.0; http://planttfdb.cbi.pku.edu/download.php) using BLASTX with E-value threshold of 10^-5^.

### Quantitative RT-PCR analysis

Total RNA was extracted by using the RNeasy Plant Mini Kit (Qiagen, Valencia, CA, USA) following the manufacturer’s instructions. The quality of the isolated RNA was checked on ethidium bromide-stained agarose gels, and its concentration was calculated according to the measured optical density (OD) of the samples at 260 and 280 nm (DropSense96C Spectrophotometer, Trinean, Belgium). The 1 μg of the total RNA was used for the cDNA synthesis using SuperScript^TM^ III first strand RT-PCR kit (Invitrogen, Carlsbad, CA, USA) with an oligo(dT)_20_ primer. After cDNA was obtained from *S. tora*, qRT-PCR was performed using gene-specific primers (S1 Table). Real-time PCR analysis was optimized and performed using the Roche LightCycler^®^ 480 II instrument and SYBR^®^ Green Real-Time PCR Master Mix (Bio-Rad, InS., Hercules, CA, USA) under condition of an initial denaturation at 95°C for 30 s followed by 40 cycles of denaturation at 95°C for 10 s, annealing and extending at 55°C for 15 s. The relative expression of specific genes was quantified using the 2^-ΔΔCt^ calculation according to the manufacturer’s software [35] (where ΔΔC_t_ is the difference in the threshold cycles), and the internal reference gene was the elongation factor 2 for data normalization. Reliability of the amplification parameters was analyzed at 1:15 dilutions of the cDNA samples. The mean threshold cycle values for the genes of interest were calculated from three experimental replicates.

### Extraction of anthraquinones and LC-MS analysis

Early- and late-stage of seed samples were extracted with methanol using sonication for 30 min at 60°C. After extraction, samples were centrifuged at 12,000 rpm for 3 min at 25°C and the supernatant was filtered with 0.2 μm Acrodisc^®^ MS Syringe Filters with WWPTFE membrane (Pall Corporation, Port Washington, NY, USA). Quantitative analysis of anthraquinones was performed by a Triple TOF 5600+ Spectrometer with a DuoSpray ion source (AB Sciex, Ontario, CA, USA) coupled with a Nexera X2 UHPLC (Shimadzu, Kyoto, Japan) equipped with binary solvent manager, sample manager, column heater, and photodiode array detector. UHPLC was performed on a ACQUITY UPLC®BEH C18 column (1.7 μm, 2.1 x 100 mm, Waters Corporation, Milford, USA) and mobile phases consisted of 5 mM ammonium acetate in water (eluent A) and 100% acetonitrile (eluent B). The gradient profile was as follows: 0-1 min, 20% B; 1-3.5 min, 10-30% B; 3.5-8 min, 30-50% B; 8-12 min, 50-100% B; 11-17 min, 100% B. The flow rate was 0.5 mL/min and five microliters of samples were injected. For detecting peaks from test samples, MS parameter in ESI-negative mode was used as follows: nebulizing gas, 50 psi; heating gas, 50 psi; curtain gas, 25 psi; desolvation temperature, 500°C; ion spray voltage floating, 4.5 kV.

### Data availability

The RNA-Seq and Iso-Seq sequences generated from Illumina and PacBio RS II sequencing of four tissue samples of *S. tora* were deposited at the National Center for Biotechnology Information (NCBI) Sequence Read Archive database with the accession number SRP159435.

## Results and discussion

### RNA sequencing and de novo transcriptome assembly

*De novo* transcriptome analysis is a good tool for generating the overall genetic information of an organism without full genome sequencing and leads to discoveries of new genes, molecular markers, and tissue-specific expression patterns. We used the Illumina HiSeq 2500 system and PacBio RS II platform to sequence the cDNA libraries of the leaf, root, and early- and late-stages of seed for elucidating secondary metabolites biosynthesis and understanding their spatial and temporal expression pattern in *S. tora*. Illumina Hiseq 2500 sequencing platform produced 278,031,495 raw reads and averaged 23,169,291 reads per tissue (S2 Table). In total, more than 270 million reads showed high quality read rates (Q30 values) of over 88.00% (S2 Table). The Trinity assembler from the four different libraries generated a total of 118,635 unigenes that were more than 300 base pairs (bp) long (Fig 1). The length of the transcripts varied from 300 to 18,622 bp with an average length of 832.25 bp, the N50 length of 1,082 bp, and the GC content of 39.51% (Table 1).

**Fig 1.**
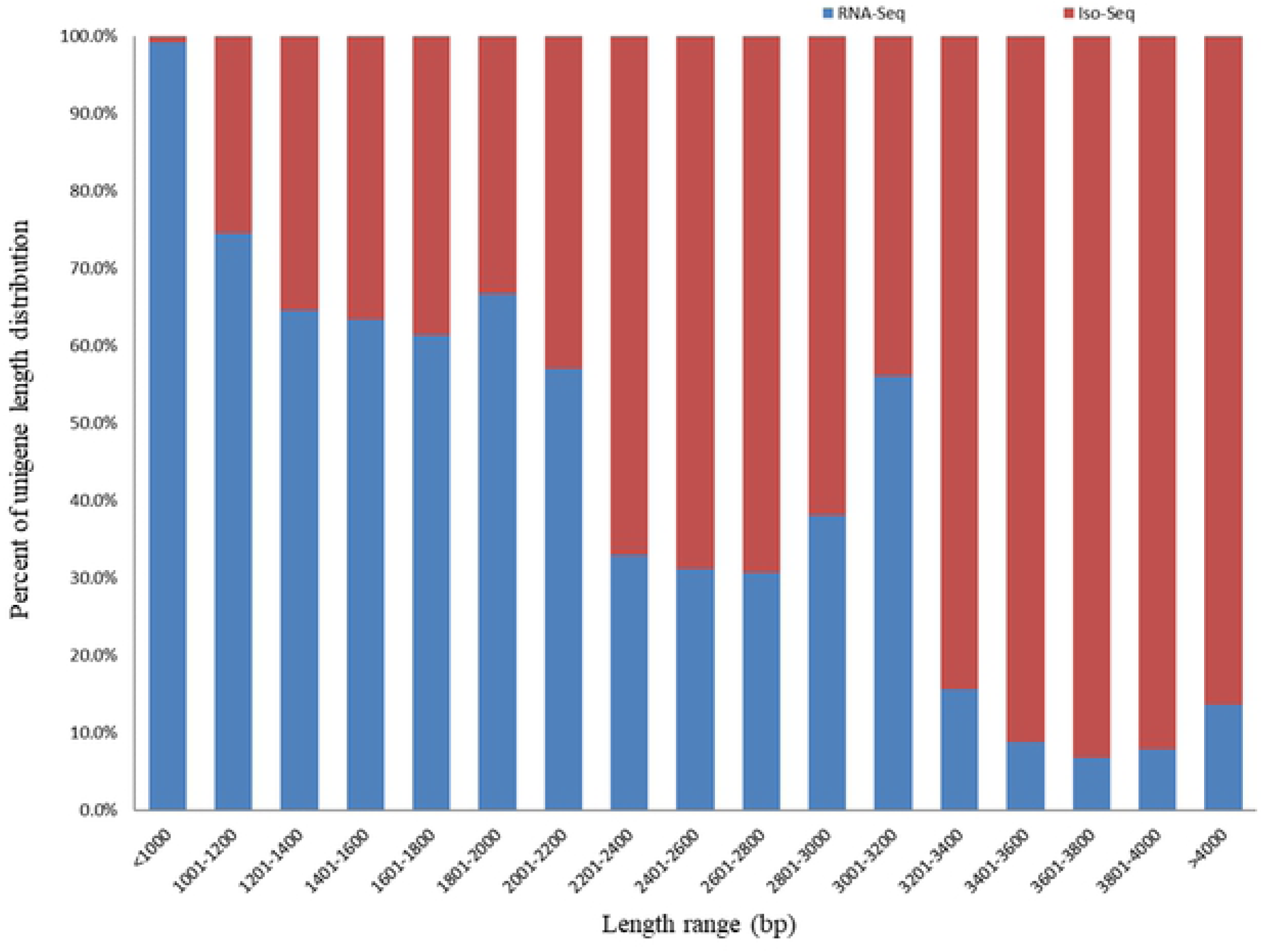
The length distribution of transcripts in *S. tora.* X and Y axis represent unigene lengths and percent of unigene length distribution, respectively.

**Table 1.** Assembly statistics of the *S. tora* transcriptome by RNA-Seq and Iso-Seq.

| Assembly statistics | RNA-Seq | Iso-Seq |
| --- | --- | --- |
| Number of unigenes | 118,635 | 39,364 |
| Total size (bp) | 98,734,027 | 112,216,332 |
| Minimum length (bp) | 300 | 435 |
| Maximum length (bp) | 18,622 | 6,814 |
| Average length (bp) | 832 | 2,851 |
| N50 length (bp) | 1,082 | 3,513 |
| GC contents (%) | 39.51 | 38.60 |

A unigene, the assembled transcript that represents a hypothetical gene, can be represented by several isomers as different forms of the same protein. The PacBio RS II sequencing platform produced 768,745 raw reads. After classification and clustering, 118,703 high-quality isoforms were obtained from three different libraries, which contained 39,672, 32,954, and 46,077 high-quality isoforms per library sizes (<2 kb, 2-3 kb, and >3 kb) (S3 Table). The 118,703 high-quality isoforms from three different libraries generated 39,364 non-redundant unigenes after the CD-HIT-EST program removed redundant isoforms. The total size of the assembly was 112 MB with 57% of transcripts larger than 500 bp and 12% larger than 2,000 bp. In total, our analysis generated two unigene sets: 118,635 from RNA-Seq and 39,364 from Iso-Seq (Fig 1). The two unigene sets showed similar GC contents. However, overall unigene lengths of each set showed that the length of the Iso-Seq was longer than RNA-Seq. Unigenes obtained from Iso-Seq were better in terms of minimum length, average length, and N50 length (Table 1).

In our analyses, we used the Iso-Seq unigene set mainly as a reference for RNA-Seq data. Due to other dissimilar characteristics, such as the transcript length between the RNA-Seq and Iso-Seq gene sets, this study did not constitute an integrated unigene set. Later, we plan to create one using the reference-guided method when the *S. tora* genome sequencing is completed.

### Functional annotation and classification

Annotation of function is required to characterize transcripts and understand the complexity and diversity of an organism. For the functional annotation, the assembled 118,635 unigenes obtained from RNA-Seq of leaf, root, early seed, and late seed tissue samples were screened using an FPKM criterion of ≥ 1, which resulted in 56,707 unigenes. To obtain the best annotations, assembled 56,707 RNA-Seq unigene sets and 39,364 Iso-Seq unigene sets of *S. tora* were aligned with four public protein databases. We used the BLASTX program against NCBI Nr, Swiss-Prot, KEGG, and GO protein databases with an E-value threshold of 1e-5. Annotations of RNA-Seq and Iso-Seq unigenes resulted in the identification of 43,286 and 36,882 unigene sets, which were respectively matched with known proteins. The Venn diagram displays the unique best BLASTX hits from NCBI Nr, Swiss-Prot, KEGG, and GO databases (S1 Fig). The overlapping regions of the four circles indicate the number of unigenes sharing BLASTX similarities in respective databases. The Venn diagram of RNA-Seq showed significant matches: 32,469 to Swiss-Prot (75.01%), 42,552 to NCBI Nr (98.30%), 3,279 to KEGG (7.58%), and 30,287 to GO terms (69.97%). So did the Venn diagram of Iso-Seq: 30,626 to Swiss-Prot (83.04%), 36,830 to NCBI Nr (99.86%), 6,441 to KEGG (17.46%), and 26,762 to GO terms (72.56%). In summary, 43,286 RNA-Seq and 36,882 Iso-Seq unigene sets had at least one significant protein match to these databases. The pattern of annotation of RNA-Seq and Iso-Seq showed that the Iso-Seq is better than RNA-Seq at annotating essential data. Non-significant genes that may represent new genes, non-coding RNA, or RNA representing unnecessary information is not evaluated in this annotaion, and further analysis is required. Matches to the Nr database also indicated that a large number of the *S. tora* unigenes closely matched the sequences of *Glycine max* (26.94%), *Glycine soja* (13.07%), *Vigna radiate* var. *radiata* (3.21%), *Cicer arietinum* (9.38%), and *Phaseolus vulgaris* (5.63%). Unigenes of 15 species in the Nr database had > 1% match with those of *S. tora* (S2 Fig).

To further functionally characterize the *S. tora* transcriptome, we classified the functions of RNA-Seq and Iso-Seq unigenes using GO analysis. The distribution of RNA-Seq and Iso-Seq unigene sets in different GO categories is shown in Fig 2. The three main categories of GO annotations of RNA-Seq included 26,616 GO terms (42.12%) for biological process, 20,211 terms (31.98%) for molecular function, and 16,365 terms (25.90%) for cellular component. Among biological process, organic substance metabolic process (17.00%) and primary metabolic process (16.00%) were the most abundant GO categories. Regarding molecular function, GO terms related to organic cyclic compound binding (19.00%) and heterocylic compound binding (19.00%) were the most abundant, while cell part (22.00%) and cell (22.00%) were the mostly represented GO categories in cellular components. Conversely, the three main categories of GO annotation of Iso-Seq include 57,137 GO terms (45.64%) for biological process, 31,562 terms (25.13%) for molecular function, and 36,876 terms (29.37%) for cellular component. Among biological process, organic substance metabolic process (16.00%) and primary metabolic process (16.00%) were the most abundant GO categories of biological process. The GO terms related to nucleotide binding (16.00%) and nucleoside phosphate binding (16.00%) were the most abundant in molecular function categories. Also, the most abundant GO categories in cellular component were cell part (24.00%) and cell (24.00%). GO terms pattern of RNA-Seq and Iso-Seq was similar in patterns.

**Fig 2.**
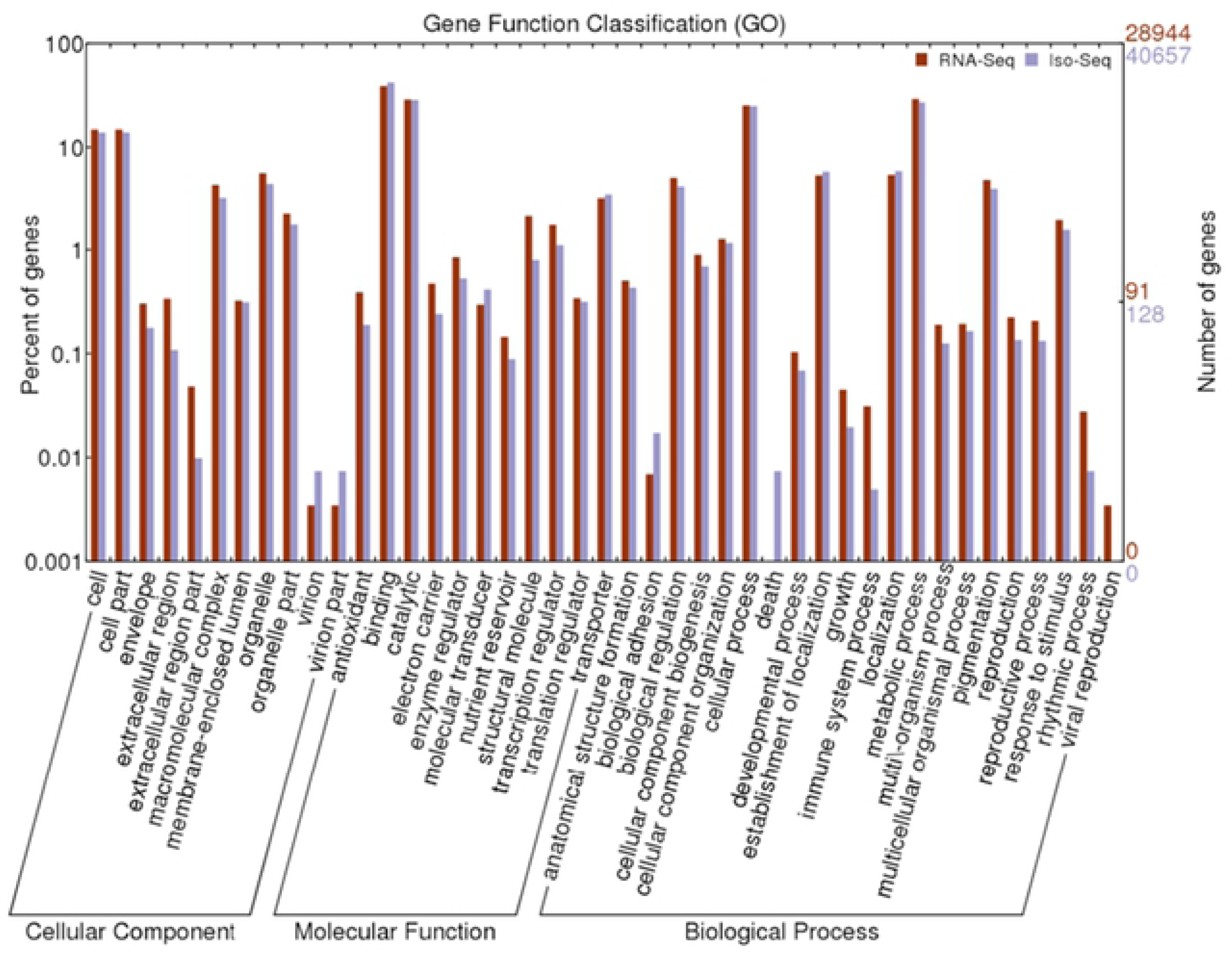
Histogram of gene ontology (GO) classification from RNA-Seq and Iso-Seq. The results are summarized in three main categories: biological process, molecular function, and cellular component.

Transcription factor (TF) families, including ARF, bHLH, bZIP, C2H2, ERF, MIKC, MYB, NAC, and WRKY, play a key regulatory role in the expression of genes, which are involved in plant secondary metabolism and response to environmental stress, by binding to specific cis-regulatory elements of the promoter regions. The number of genes encoding for different TF families varies in different plants to perform species-/tissue-specific or developmental stage-specific function [36]. In our study, 3,284 RNA-Seq and 3,576 Iso-Seq were generated with a total of 6,860 unigenes assigned to 56 TF families. Among these, bHLH (521, 15.86%) were found to be the most abundant in RNA-Seq followed by WRKY (243, 7.40%), C2H2 (189, 5.76%), MYB (177, 5.39%), bZTP (170, 5.18%), and NAC (150, 4.57%). Similarly, in the Iso-Seq, bHLH were found to be the most abundant followed by WRKY, but the other TF families showed a slight ranking change (S3A Fig).

Expression of the gene varies depending on the environment in which each species is exposed, and specific or large amounts of the gene are expressed. The degree of expression of the TF family, which mediates and controls their expression, is essential for the molecular genetics of organisms, so in order to investigate tissue specific gene expression in *S. tora* we studied the expression of genes in leaf, root and early and late seed tissues. Interestingly, different expression patterns for TFs were observed in four tissues of *S. tora*. Some TFs were unique to each tissue, whereas others were enriched in respective tissues. The 35 and 98 TFs among a total of 133 TFs expressed in leaf, 41 and 97 from 138 TFs in root, 30 and 51 from 81 TFs in early-stage, and 15 and 18 from 33 TFs in late-stage during seed development were tissue-enriched and -specific (S3B Fig). Notably, growth regulating factor (GRF) in the TF family was dominantly expressed in late-stage seed tissue (S4 Table). GRFs are plant-specific transcription factors that were originally identified for their roles in stem and leaf development [37]. However, recent studies highlight its importance in other central developmental processes including root development, flower, and seed formation. Expression of GRFs has also been observed in various rice and maize tissues, suggesting their involvement in seed development [38, 39].

### Differential gene expression analysis during seed development

To compare genes of *S. tora* with differential expression level in late-stage seed development to those in early-stage development, we used the DESeq method. The transcripts with log2 fold change (FC) >1 and p-value < 1e-3 were considered as differentially expressed genes (DEGs). Pair-wise comparison of transcripts between early- and late-stages of seed development resulted in a total of 14,825 DEGs in RNA-Seq. As seeds matured, 4,935 genes were identified as up-regulated and 9,890 genes were down-regulated. These genes belong to diverse functional groups including glycosyl hydrolases, dehydrogenases, transferases, kinases, phosphatases, cytochrome P450, oxygenases, and hormone-responsive proteins. A heat map was constructed to cluster the top 50 DEGs based on the similarity and diversity of expression profiles using normalized FPKM values within a range of 6 to 16 (Fig 3). Specifically, transcripts of various proteins are expressed differently depending on the tissue and stage of seed. In early-stage seeds, the expression of chalcone synthase, peroxidase, and cell wall/vacuolar inhibitor of fructosidase were higher than those of late-stage seeds. In particular, C/VIF releases glucose and fructose in irreversible reactions, which is essential to plant growth, storage compound accumulation, and stress response [40]. Conversely, in late-seed development, late embryogenesis-abundant (LEA) proteins and heat shock proteins (HSPs) appeared to be more abundant than early seeds like the adlay species [41].

**Fig 3.**
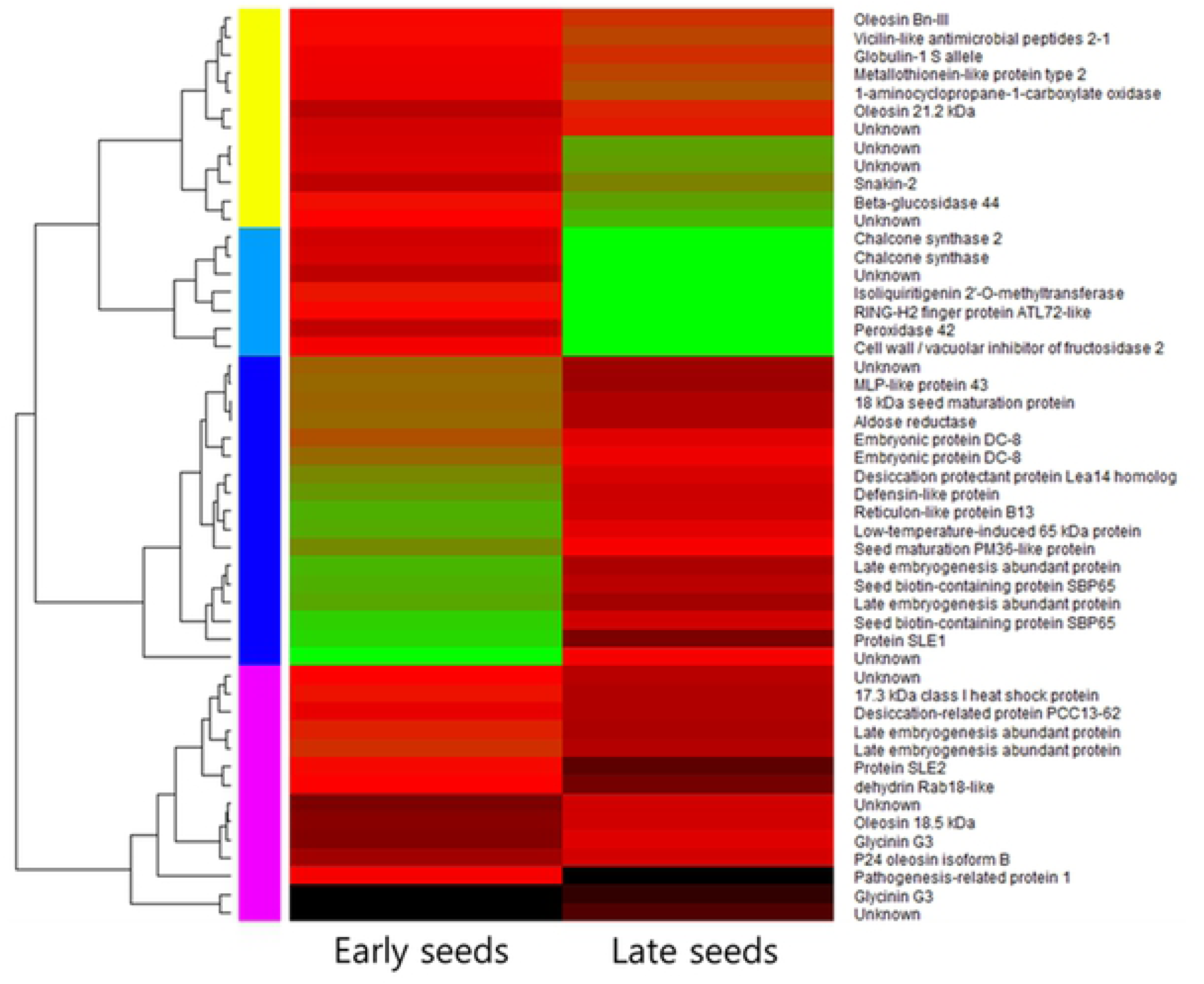
Heat map of top 50 differentially expressed genes between early- and late-stages of seed development in *S. tora.* Heatmap showing differentially expressed genes between early and late stages of seed development in *S. tora*. Color scale representing normalized expression values is shown at the bottom.

Previously, the expression of genes in leaves, roots, and early- and late-seed tissues were examined to investigate the tissue-specific gene expression of *S. tora.* During this process the transcripts exhibiting tissue-specific expression were identified and the top 10 transcripts were selected (S4 Fig). Real-time PCR analysis was performed in order to accurately identify differential expression of selected transcripts in the data. Expression analysis was carried out from the selected genes belonging to carbohydrate mechanism, the secondary metabolite pathway, and the associated transcription factors (Fig 4). These results were consistent with tissue-specific gene expression data in various tissues. As results, 3 genes were identified in the qRT-PCR of the seeds to be specifically expressed compared to other tissues. Cell wall/vacuolar inhibitor of fructosidase 2(C/VIF2) play important roles in carbohydrate metabolism, stress responses, and sugar signaling. The specific expression of C/VIF2 in early seeds is implicated in several mechanisms of maturation. Cytochrome P450 83B1 genes showed the highest expression levels in leaf, followed by root, late seed, and early seed. Cytochrome P450 83B1 protein is known to be involved in auxin homeostasis and glucosinolate biosynthesis associated with plant growth and pathogenic responds [42]. Also, seed biotin-containing protein gene showed the highest expression levels in late seed, followed by early seed, demonstrating that the protein plays an important role in the developmental stage of the seed. And organic-cation/carnitine transporter 1 protein gene expressed high levels in root, followed by leaf and late seed. Organic-cation/carnitine transporter families are generally characterized as polyspecific transporters involved in the homeostasis of solutes in animals [43]. Although some publications have suggested that this protein is known as stress-regulated member of plants and that it is involved in plant growth [43], little is known about the function, localization, and regulation of plants.

**Fig 4.**
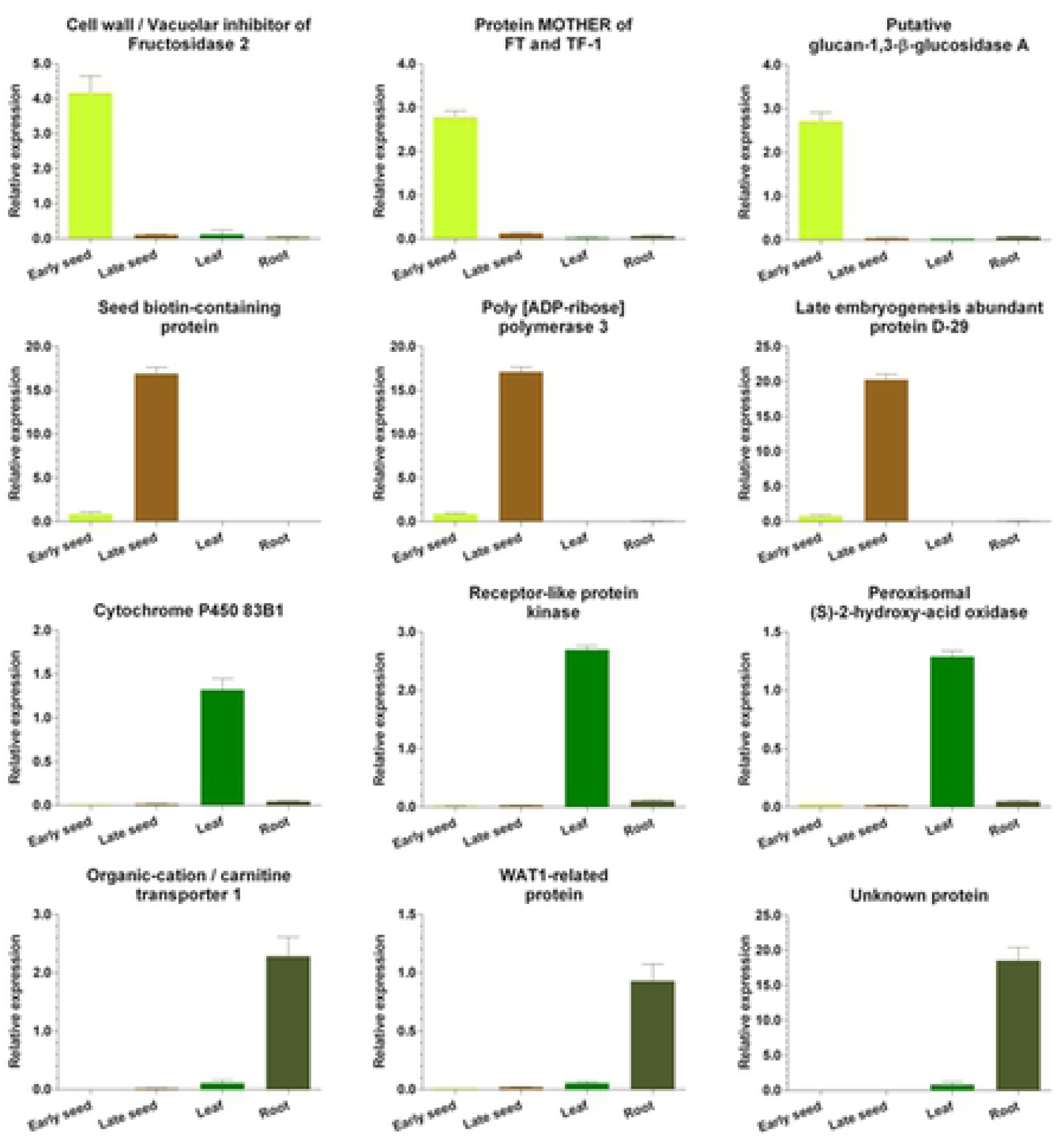
Real-time PCR validation of gene expression obtained via RNA-Seq. All the real-time PCR experiments were performed at least three times in each independent biological experiment (3 replicates). Error bars represent SEM from triplicates.

To determine the biological function of DEGs during seed development, GO classification analysis was carried out using Blast2GO. The results showed that 25 functional groups, including 3 major ontologies, classified 63,192 GO terms annotated by the GO database: biological process, cellular component, and molecular function. Many of these DEGs were dominant catalytic activity, binding metabolism, cellular processes, cell parts and cells (S5 Fig). In confirming whether there is specificity for development of seeds in relation to their transcripts, orthologous *S. tora* genes were applied to gene ontology enrichment analysis using the AgriGO program. In molecular function of GO ontologies, the level of binding function was increased in the up-regulated DEGs. Among them, RNA binding increased to a very high level. In addition, down-regulated DEGs showed an increase in the catalytic activity function, and they also increased protein kinase activity, transferase activity, and microtubule motor activity (S5 Fig).

To identify specific metabolic pathways that are responsible for the transcriptional changes of enzymatic genes during seed development of *S. tora*, we performed MapMan analysis with the expression data of genes showing at least 2-fold differential expression between seed developmental stages. We made the figure to depict the biological processes of interest, and display log2-normalized expression counts onto pictorial diagrams. Most of the genes in cell metabolism are involved in cell wall metabolism, lipid metabolism, carbohydrate metabolism, and secondary metabolism. The dynamic changes in metabolic pathways during seed development were provided in Fig 5, in which we identified the downward trend of overall transcription in the seed development process. In particular, it was clear that lipid metabolism, precursor synthesis, flavonoid metabolisms, and phenylpropanoids/phenolics metabolisms were down-regulated, while the FA synthesis of lipid metabolism and the N-msc of secondary metabolism were up-regulated.

**Fig 5.**
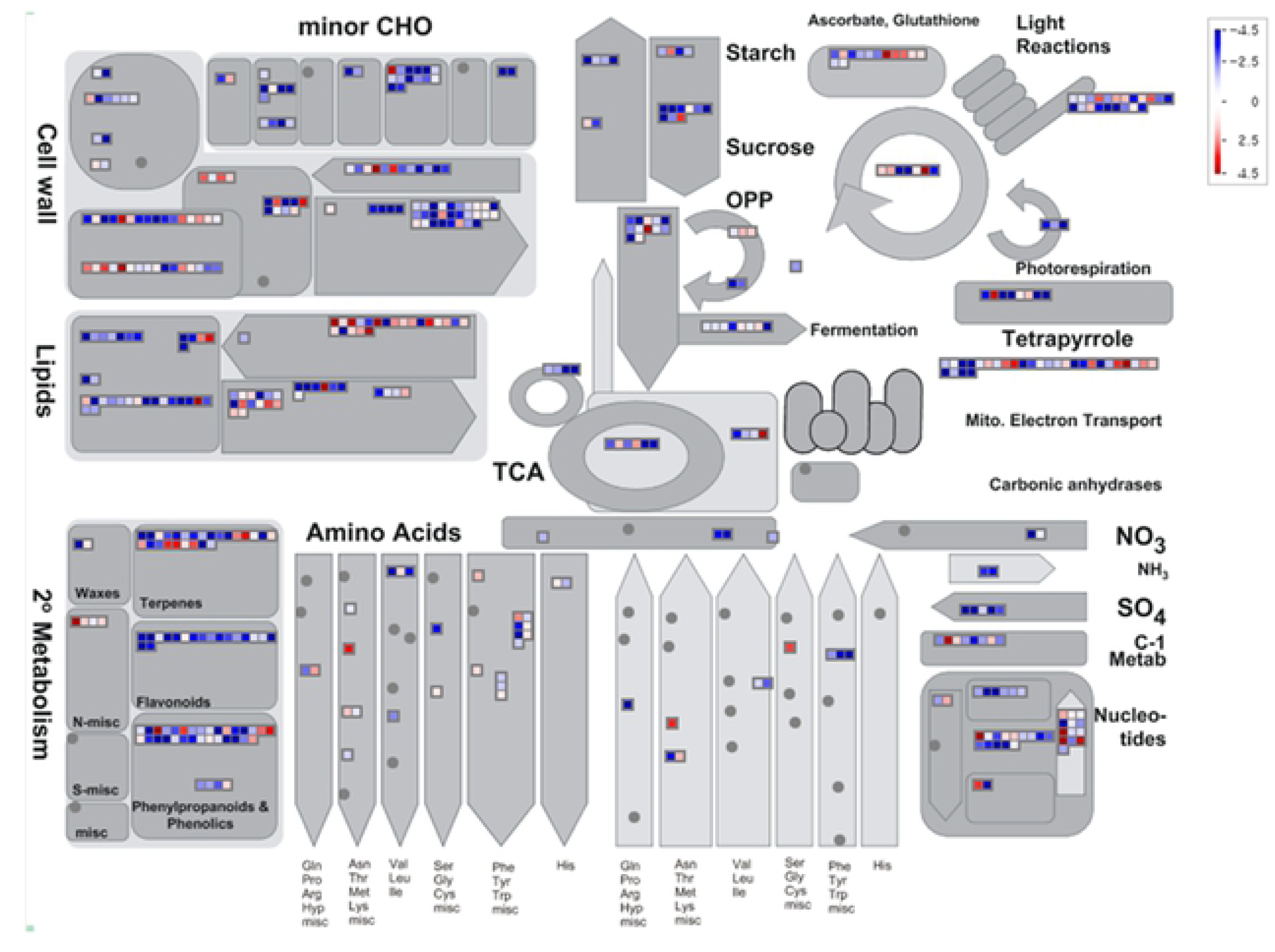
MapMan metabolism overview maps showing differences in transcript levels during seed development. MapMan software was used to provide a snapshot of modulated genes over the main metabolic pathways. Log2 fold changes values are represented. Up-regulated and down-regulated transcripts are shown in red and blue, respectively.

### Candidate gene families involved in anthraquinones biosynthesis

*S. tora* is well known for its various therapeutic effects (e.g., for its anti-hypertensive, diuretic, anti-cancer, anti-microbial and cholesterol-lowering effects). Each effect is caused by various secondary metabolites produced in *S. tora*, the best known of these being anthraquinone. The biosynthesis of anthraquinone shares isochorismate pathways with phenylpropanoid and shares MEP/DOXP, MEV, and shikimate pathways with carotenoid and flavonoid. In addition, the polyketide pathway is an important part of the anthraquinone biosynthesis. To analyze the active biosynthesis of anthraquinones, we determined the contents of seven compounds of the anthraquinone biosynthesis pathway in early- and late-seed tissues. As seeds matured, anthraquinone compounds were more accumulated in late seed than early seed (Fig 6 and Table 2). Among the seven compounds, gluco-obtusifolin has the highest content in seed tissues (Fig 6 and Table 2). It is well known that aurantio-obtusin is the most significant active compound [8] and is distributed mainly in the seed [44]. However, we found that low levels of aurantio-obtusin were observed at the early and late developmental stages. A possible explanation for this reason is that aurantio-obtusin may accumulate mainly in the matured and/or dry seed.

**Fig 6.**
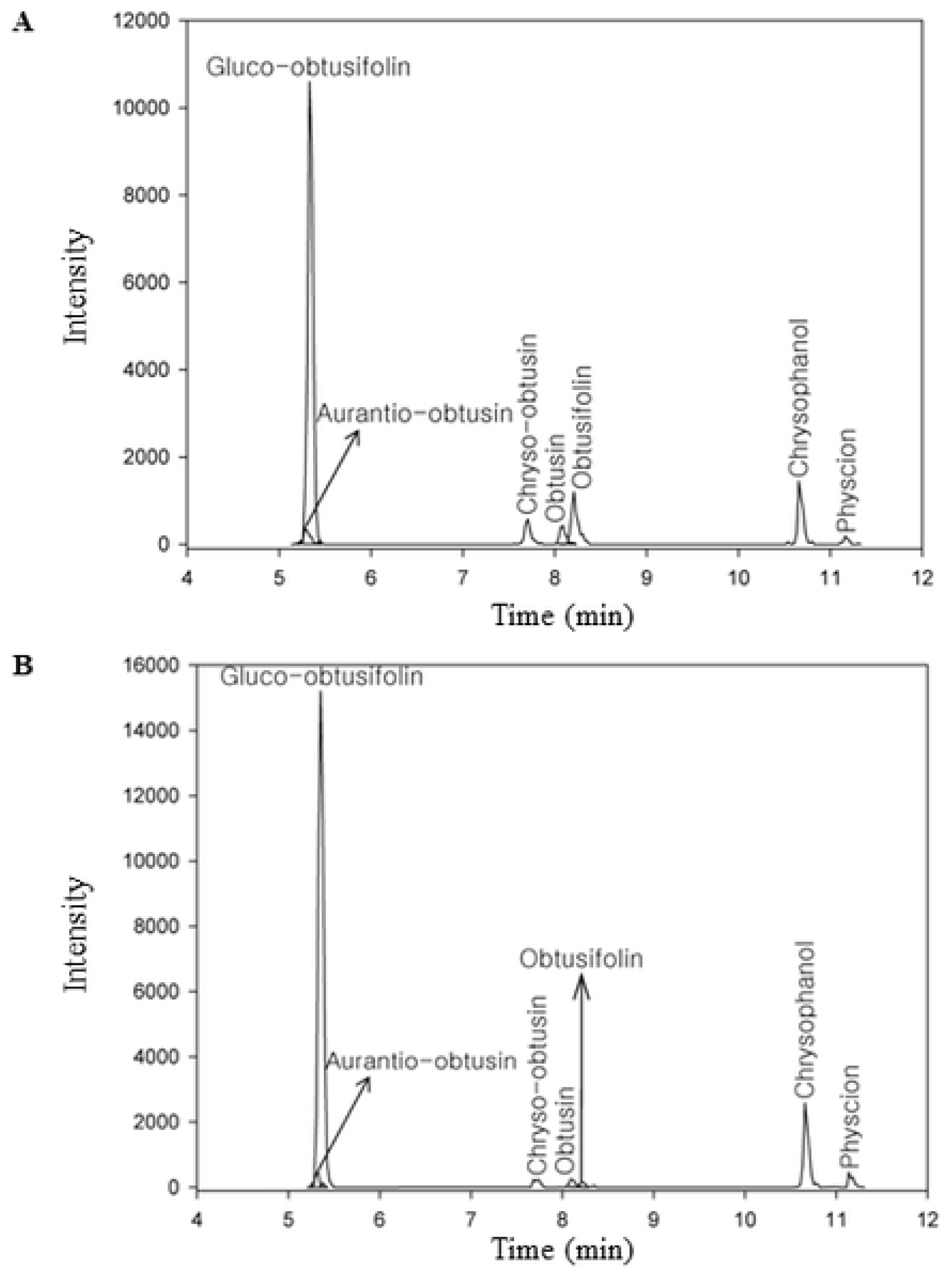
GC-MS analysis of anthraquinone during seed development. Seven anthraquinone levels in the early seed (A) and in the late seed (B).

**Table 2.** Anthraquinone contents in the early and late seeds.

| Compounds | Formula | RT<sup>a</sup> | Contents (ug/g) |  |
| --- | --- | --- | --- | --- |
|  |  |  | Early Seed | Late Seed |
| Gluco-obtusifolin | C22H22O10 | 5.31 | 80.72<sup>b</sup> | 141.27 |
| Aurantio-obtusin | C17H14O7 | 5.27 | 1.07 | 1.08 |
| Chryso-obtusin | C19H18O7 | 7.68 | 1.39 | 0.69 |
| Obtusin | C18H16O7 | 8.07 | 0.83 | 0.60 |
| Obtusifolin | C16H12O5 | 8.19 | 1.38 | 0.26 |
| Chrysophanol | C15H10O4 | 10.65 | 11.39 | 6.69 |
| Physcion | C16H12O5 | 11.14 | 2.02 | 0.96 |
| Total | - | - | 98.80 | 151.55 |

<sup>a</sup> indicates retention time.
<sup>b</sup> represents mean of three replicate experiments.

To observe gene expression levels of each parts and to compare the changes in gene expression levels between different parts, their levels were normalized to the FPKM (reads per kilobase of exon model per million mapped reads), and transcripts were hierarchically clustered based on the Log2(FPKM+1), allowing us to observe the overall gene expression pattern (Fig 7). In our study, there were 337 RNA-Seq and 212 Iso-Seq genes involved in *S. tora* secondary metabolites, and they were classified into five pathways including the MEP/DOXP, MEV, shikimate, carotenoid, and flavonoid/polyketide (Fig 7 and S5 Table). There were 35 RNA-Seq and 24 Iso-Seq genes in *S. tora* for seven enzymes involved in MEP/DOXP pathway and mevalonate pathway leading to production of precursor dimethylallyl disphosphate (Fig 7 and S5 Table). They are also involved in the shikimate pathway leading to the production of precursor 1,4-dihydroxy-2-napthoyl-CoA including 40 RNA-Seq and 31 Iso-Seq genes for 9 enzymes (DAHPS, DHQS, DHQD/SDH, SMK, EPSP, CS, ICS, MenE, and MenB). In MEP/DOXP, 13 DXPS (1-deoxy-_D_-xylulose-5-phosphate synthase, EC 2.2.1.7) were expressed in anthraquinone synthesis. In them, DN49358_C0_g1 was expressed in large amounts up to the early stage of seed, but appeared to be greatly reduced by the late stage. This gene was also expressed at high levels in leaf and root tissues. Furthermore, DN27315_c0_g1 demonstrated higher levels of gene expression in leaf than in other tissues. And only three of the 13 DXPS genes showed high levels of expression independent of tissue and seed development. ISPD, CDPMEK, and ISPF genes were identified in only 1 and 2, while HDS and HDR were identified in more frequent. HDS and HDR were identified in genes 8 and 6, and HDS ((E)-4-hydroxy-3-methylbut-2-enyl-diphosphate synthase, DN48094_c1_g1) and HDR (4-hydroxy-3-methylbut-2-enyl-diphosphate reductase, DN25595_c0_g1) showed high levels of expression regardless of tissue and seed development. In the MEV pathway, ACCA (acetyl-CoA carboxylase) was identified in 29 genes, and 3 genes (DN51063_c1_g1, DN51063_c2_g1, and DN72707_c0_g1) sustained high levels of expression independent of tissue and seed development. Conversely, one HMGR (DN9882_c0_g1) was down-stream of expression level. Except for some genes, ACCA, HMGS, HMGR, MK, PMK, and MPD of expression levels are down-stream, and 1 of 4 IPPS (isopentenyl-diphosphate delta-isomerase, DN67602_c1_g1) genes showed high level of expression independent of tissue and seed development.

**Fig 7.**
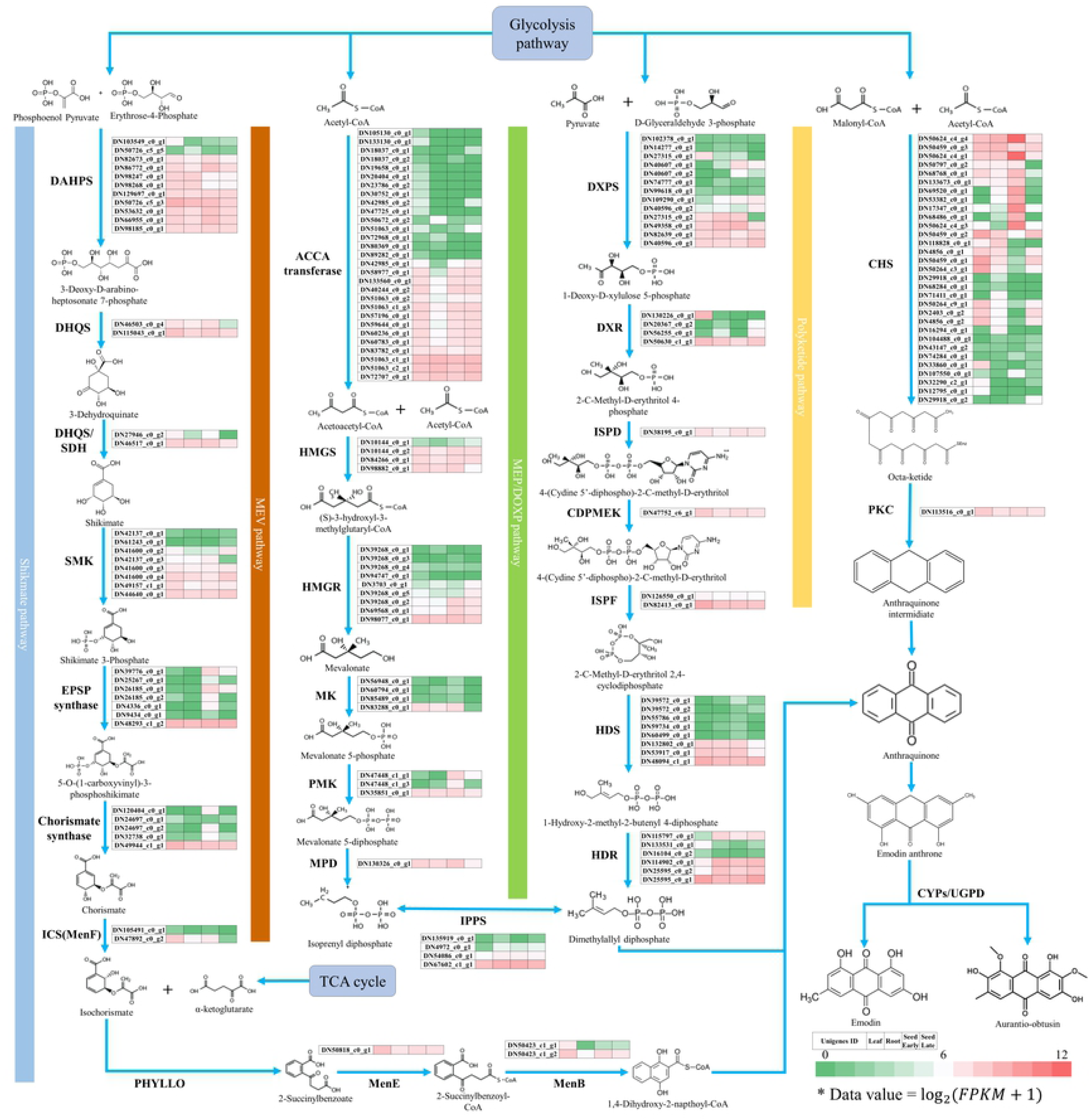
The up-down of putative genes of anthraquinone-biosynthetic pathway in *S. tora.* It was normalized to the FPKM to compare the changes in gene expression levels between different parts of *S. tora*. Total gene expression levels were clustered based on the Log2 (FPKM +1). DXPS, 1-Deoxy-_D_-xylulose-5-phosphate synthase (EC 2.2.1.7); DXR, 1-Deoxy-_D_-xylulose-5-phosphate reductoisomerase (EC 1.1.1.267); ISPD, 2-C-Methyl-_D_-erythritol 4-phosphate cytidylyltransferase (EC 2.7.7.60); CDPMEK, 4-Diphosphocytidyl-2-C-methyl-_D_-erythritol kinase (EC 2.7.1.148); ISPF, 2-C-Methyl-_D_-erythritol 2,4-cyclodiphosphate Synthase (EC 4.6.1.12); HDS, (E)-4-Hydroxy-3-methylbut-2-enyl-diphosphate synthase (EC 1.17.7.1); HDR, 4-Hydroxy-3-methylbut-2-enyl diphosphate reductase (EC 1.17.1.2); ACCA, Acetyl-CoA carboxylase (EC 6.4.1.2); HMGS, Hydroxymethylglutaryl-CoA synthase (EC 2.3.3.10); HMGR, Hydroxymethylglutaryl-CoA reductase (EC 1.1.1.34); MK, Mevalonate kinase (EC 2.7.1.36); PMK, Phosphomevalonate kinase (EC 2.7.4.2); MPD, Methyl parathion hydrolase (EC 3.1.8.1); IPPS, Isopentenyl-diphosphate delta-isomerase (EC 5.3.3.2); DAHPS, 3-Deoxy-7-phosphoheptulonate synthase (EC 2.5.1.54); DHQS, 3-Dehydroquinate synthase (EC 4.2.3.4); DHQD/SDH, 3-Dehydroquinate dehydratase/shikimate dehydrogenase (EC 4.2.1.10/1.1.1.25); SMK, Shikimate kinase (EC 2.7.1.71); EPSP, 3-Phosphoshikimate 1-carboxyvinyltransferase (EC 2.5.1.19); CS, Chorismate synthase (EC 4.2.3.5); ICS, Isochorismate synthase (EC 5.4.4.2); PHYLLO, 2-Succinyl-5-enolpyruvyl-6-hydroxy-3-cyclohexene-1-carboxylic acid synthase (EC 2.2.1.9); MenE, 2-Succinylbenzoate-CoA ligase (EC 6.2.1.26); MenB, 1,4-Dihydroxy-2-naphthoyl-CoA synthase (EC 4.1.3.36); GGPS, Geranylgeranyl diphosphate synthase (EC 2.5.1.1); PSY, Phytoene synthase (EC 2.5.1.32); PDS, Phytoene desaturase (EC 1.3.99.30); ZDS, Zeta-carotene desaturase (EC 1.3.5.6); LYCB, Lycopene beta-cyclase (EC 5.5.1.19); LYCE, Lycopene epsilon-cyclase (EC 5.5.1.18); BCH, Beta-carotene hydroxylase (EC 1.14.13.129); ZEP, Zeaxanthin epoxidase (EC 1.14.15.21); PAL, Phenylalanine ammonia-lyase (EC 4.3.1.24); C4H, Cinnamate-4-hydroxylase (EC 1.14.13.11); 4CL, 4-Coumarate-CoA ligase (EC 6.2.1.12); and CHS, Chalcone synthase (EC 2.3.1.74).

Anthraquinones are also known to be produced from acetyl-CoA and malonyl-CoA through polyketide pathway in plants. Chalcone synthase (CHS), a type III polyketide synthase, is an important enzyme involved in the polyketide pathway [45]. We have identified 27 RNA-Seq and 23 Iso-Seq genes encoding for enzyme involved in type III polyketide synthase (S5 Table). As a ubiquitous enzyme in higher plants, CHS is known to produce flavonoids by catalyzing the sequential decarboxylative reaction with 3 malonyl-CoA and p-coumaroyl-CoA as a starter and extender unit, respectively [46]. It was also suggested that polyketide synthase could form an anthraquinone precursor using acetyl-CoA and malonyl-CoA. And the formed precursor, octaketide is cyclized by PKC-encoding polyketide cyclase, and usually forms three-ring structures named A, B, and C rings [47]. The formed intermediate is modified by P450 to produce anthraquinone or emodin anthrone, and also to produce sennoside by modification of glycosyltransferases. These 27 PKS gene sizes averaged 584.03 bp, and the longest was 1,580 bp. Among them, only 3 genes (DN50459_c0_g1, DN2403_c0_g2, and DN50459_c0_g2) showed high levels of expression change in seed development. It seems that these genes are changing a lot in order to make the backbones of the flavonoid and carotenoid components needed for survival in the later stages of seed development. In particular, 5 genes (DN17347_c0_g1, DN50624_c4_g3, DN69520_c0_g1, DN50624_c4_g1, and DN50624_c4_g4) showed a large amount of expression in the early part of the seed, whereas in the latter part, the level of expression decreased sharply, suggesting that those genes play a very important role in the biosynthesis of the backbone of the material needed in early seed development.

In general, glycosylation is carried out at the end of secondary metabolites biosynthesis and improve the solubility and stability of the secondary metabolites. In nature, UDP-glycosyltransferases (UGT) normally facilitates glycosylation, and makes the natural product with glucose at the hydroxyl group [48]. In our study, there were 59 genes in seed stage of *S. tora*. Based on the results, 33 out of 59 genes showed more expression at the late-seed than at the early-seed stage, whereas 26 showed more expression at the early-seed stage (Fig 7 and S5 Table). The degree of expression of the seven genes (DN131354_c0_g1, DN67413_c0_g1, DN49988_c0_g2, DN50503_c0_g2, DN82643_c0_g1, DN17331_c0_g2, and DN137099_c0_g1) seems to increase rapidly during the growth of the seed, which seems to be necessary for the process of stockpiling the energy required for seed germination. In addition, DN17331_c0_g2 and DN82643_c0_g1 seem to have a great effect on the glycosylation during seed development because they undergo a significant amount of change. Conversely, the expression level of the four genes (DN50189_c2_g1, DN11235_c0_g1, DN62590_c0_g1, and DN76515_c0_g1) seemed to decrease rapidly, and the remaining 22 genes were found to be expressed with a relatively small decrease.

## Conflict of interest

The authors declare that they have no conflict of interest.

## Acknowledgements

This work was carried out with the support of National Institute of Agricultural Sciences [Project no. PJ013818] and Cooperative Research Program for Agriculture Science and Technology Development [Project title: National Agricultural Genome Program, Project no. PJ010457], Rural Development Administration, Republic of Korea.

## Author Contributions

**Funding acquisition:** Sang-Ho Kang

**Data curation:** Joon-Soo Sim, Chang-Muk Lee, So-Ra Han

**Methodology:** So Youn Won, Soo-Jin Kwon, Jung Sun Kim

**Writing – original draft:** Sang-Ho Kang, Woo-Haeng Lee

**Writing – review & editing:** Sang-Ho Kang, Chang-Kug Kim, Tae-Jin Oh

## Supporting information

**S1 Table. Gene-specific primers used for tissue-specific qRT-PCR.**

**S2 Table. General properties of the reads produced by Illumina Hiseq 2500 sequencing platform.**

**S3 Table. General properties of the reads produced by PacBio sequencing platform.**

**S4 Table. Tissue-enriched and specific transcription factors (TFs) distribution of each tissue.**

**S5 Table. Gene associated with the secondary metabolite pathway in *S. tora*.**

**S1 Fig. The distribution of annotated unigenes by various public protein databases.** Venn diagram showing the proportion of annotated unigenes in NCBI Nr, KEGG, Swiss-Prot, and GO databases with RNA-Seq (A) and Iso-Seq (B).

**S2 Fig. Species distribution of the top BLAST hits.** Top-hit species from RNA-Seq and Iso-Seq were calculated based on sequence alignments with the lowest E-value obtained from BLAST.

**S3 Fig. Distribution of TF families of *S. tora.*** Distribution of transcripts (3,284 for RNA-Seq and 3,576 for Iso-Seq) that encode for transcription factors (A). Number of transcripts exhibiting specific expression in different tissues has been indicated by bar and table (B). Tissue-specific shows 10-fold higher FPKM in one tissue compared with three tissues, and tissue-enriched represents 5-fold higher FPKM compared with other tissues.

**S4 Fig. Heatmaps representing the top 10 genes that showed tissue-specific expression in the *S. tora* leaf, root, and early and late seeds.** Red represents high abundance and green represents low abundance.

**S5 Fig. AgriGo analysis of upregulated and downregulated genes during seed development.** A total of 4,935 (up-regulated, **A**) and 9,890 (down-regulated, **B**) genes with Molecular terms are represented by increasingly red colors. GO term enrichment was performed using single enrichment analysis (SEA) tool on AgriGo (http://bioinfo.cau.edu.cn/agrigo/). Box colors indicates levels of statistical significance: yellow=0.05; orange=e-05; and red=e-09.

